# Mapping the Causal Roles of Non-Primary Motor Areas in Human Reach Planning and Execution

**DOI:** 10.1101/2025.09.11.671853

**Authors:** Golnaz Haddadshargh, Roberto M. de Freitas, Jennifer Mak, Amy Boos, Xiaoqi Fang, Jennifer L. Collinger, Gina McKernan, Liang Zhan, Fang Liu, George F. Wittenberg

**Affiliations:** Rehab Neural Engineering Labs, University of Pittsburgh, Pittsburgh, PA; Department of Bioengineering, University of Pittsburgh, Pittsburgh, PA; Center for Neural Basis of Cognition, University of Pittsburgh and Carnegie Mellon University, Pittsburgh, PA; Department of Neurological Surgery, University of Pittsburgh 675018v1; Department of Neurology, University of Pittsburgh, Pittsburgh, PA; TECH-GRECC & HERL, VA Pittsburgh Healthcare System, Pittsburgh, PA; Department of Physical Medicine and Rehabilitation, University of Pittsburgh, Pittsburgh, PA; Department of Electrical and Computer Engineering, University of Pittsburgh, Pittsburgh, PA

**Keywords:** Brain networks, Functional brain imaging, Motor control, Reaching, Transcranial magnetic stimulation

## Abstract

Non-primary motor areas, including the dorsal premotor cortex (PMd), ventral premotor cortex (PMv), and posterior parietal cortex (PPC), are involved in movement planning, but their precise contributions remain unclear. We used functional magnetic resonance imaging (fMRI), transcranial magnetic stimulation (TMS), and kinematic assessments to explore how these areas influence reaching performance in neurologically intact adults. Participants performed a reaching task using the KINARM exoskeleton robot. Brief TMS pulse trains were initiated before movement onset to perturb cortical activity at subthreshold and suprathreshold intensities targeting bilateral PMd, PMv, and PPC. Subthreshold stimulation of contralateral PMd and PMv reduced reach efficiency and smoothness, while suprathreshold stimulation of contralateral PPC increased deviation error and reduced smoothness. Among ipsilateral targets, PMd showed consistent TMS-induced effects, and was the only target where resting-state connectivity predicted behavioral response. Stronger interhemispheric connectivity in the primary motor cortex and weaker interhemispheric PPC connectivity were associated with greater reductions in straightness and smoothness during subthreshold ipsilateral PMd stimulation. These findings show that perturbation of premotor and parietal areas during early movement execution alters reach trajectories, and differences in network organization explain variability in behavioral response. Identifying contributions of cortical areas and connectivity patterns may help personalize interventions after stroke.

This study was registered at ClinicalTrials.gov under ID NCT04286516.

## Introduction

Stroke is a leading cause of long-term disability partly because of upper limb motor impairments that impact the quality of life (Tengs and Lin 2003). Although some spontaneous recovery occurs, after a nadir in function, within the first few weeks after stroke, recovery is variable and often incomplete, particularly among individuals with severe motor deficits (Cramer 2008). In such cases, fewer than 62% regain any hand dexterity within six months of stroke onset (Kwakkel et al. 2003). Reaching and arm function also remain impaired, with movements becoming slower, more segmented, and less coordinated, and frequently relying on compensatory trunk involvement (Cirstea and Levin 2000).

Understanding the neural mechanisms supporting movement planning and execution in neurologically intact individuals provides an essential baseline for interpreting post-stroke recovery, which depends on both structural integrity and functional plasticity (Nudo 2003). Hemiparetic stroke usually involves disruption of the descending fibers of the corticospinal tract (CST), the primary pathway that mediates transmission of descending motor commands for controlling voluntary movements. This damage is a major contributor to post-stroke motor impairments, and patients with more preserved CST pathways tend to have better function from the acute through chronic stages (Stinear et al. 2007; Feng et al. 2015). The majority of CST fibers originate from the primary motor cortex (M1), but substantial projections also arise from non-primary motor areas, including the dorsal premotor cortex (PMd, ≈ 16%), ventral premotor cortex (PMv, ≈ 11%), supplementary motor area (SMA, ≈ 5%), and somatosensory cortex (Usuda et al. 2022). Damage to alternate motor pathways has also been identified as a marker of motor impairment and recovery, highlighting the role of a broader structural network in supporting functional outcomes (Lindenberg et al. 2010).

The mechanisms of motor recovery in humans remain unclear, but animal models have demonstrated functional plasticity within cortical areas underlying movement planning and execution (Schaechter et al. 2002; Inoue and Ueno 2025). PMd plays a central role in defining movement parameters, such as direction and timing, and supports externally guided movement planning (Chouinard and Paus 2006; Hoshi and Tanji 2007). Structural and functional connectivity between PMd and M1 is positively correlated with motor recovery after stroke (Schulz, Braass, et al. 2015). PMv is more directly involved in grasp-related movements, and may support recovery, particularly in cases of more extensive CST damage (Hoshi and Tanji 2007; Schulz et al. 2017). The posterior parietal cortex (PPC) has an effective role in sensorimotor integration and spatial processing of goal-directed actions (Desmurget et al. 1999). In stroke, the integrity of frontoparietal tracts has been associated with post-stroke motor performance (Schulz, Koch, et al. 2015). SMA coordinates the sequence and initiation of internally driven movements and may show increased involvement as part of compensatory adaptation (Liu et al. 2020). Despite these findings, there are remaining questions regarding the role of non-primary motor cortical areas in motor planning, and how these roles are altered after stroke.

To better understand the functional roles of non-primary motor areas in movement, researchers have increasingly used transcranial magnetic stimulation (TMS). This method allows for temporally precise imposition of neural activity that can be used to assess how a region influences a specific behavior (Paus 2005; Sack 2006). For example, short bursts of repetitive TMS (rTMS) applied over the PPC during movement initiation have been used to demonstrate its involvement in early stages of action planning (Glover et al. 2005). One limitation of the technique is limited depth of penetration and therefore selectivity for deeper structures. The medial location, depth, and anatomical variability of the SMA, for instance, limit the ability to precisely and effectively target with TMS (Immisch et al. 2001). But understanding the functional contributions of more TMS-accessible regions in intact individuals, can establish a foundation for comparing their roles after stroke and potentially guide therapeutic stimulation strategies that target preserved circuits.

Although areas like PMd and PPC have been implicated in movement planning, most studies have investigated them individually or focused on limited behavioral outcomes, leaving it unclear how different non-primary areas contribute to overall movement execution. Furthermore, the relationship between individual variability in functional connectivity and behavioral response to stimulation is not well understood. In this study we address these gaps by investigating how perturbation of different non-primary motor areas affects visually-guided reaching performance. A cohort of neurologically intact participants performed a robot-assisted reaching task while receiving TMS to six lateral frontal and parietal targets (in randomized order), and we evaluated changes in reach trajectory across stimulation conditions. Resting-state functional magnetic resonance imaging (fMRI) was used to assess whether individual connectivity differences among motor-related areas could predict stimulation-induced behavioral changes.

## Materials and Methods

### Participants

Study protocols were approved by the University of Pittsburgh Institutional Review Board (study #19080097) and registered at ClinicalTrials.gov (NCT04286516). 26 neurologically intact volunteers (16 females, 10 males; 24 right-hand dominant, 2 left-hand dominant; mean age ± standard deviation: 63.6 ± 6.8 years; age range: 47–79 years) were recruited from the Veterans Affairs Pittsburgh Healthcare System and UPMC Healthcare Systems. All participants provided written informed consent before enrollment.

Exclusion criteria included inability to provide informed consent, serious medical illness that would preclude participation, orthopedic problems limiting range of motion in the arm assessed in the study or other impairments that could interfere with study activities, visual loss sufficient to prevent viewing the test patterns on the robot computer monitor, concurrent enrollment in another greater-than-minimal-risk study, medical conditions or implants that prevent safe administration of TMS or MRI, and pregnancy.

Of the 26 recruited participants, 20 provided complete datasets for kinematic, resting-state fMRI, and task-based fMRI analyses. Six participants were excluded from the kinematic assessments: three due to protocol differences, two withdrawn after MRI scanning (one upon request and one due to poor scan quality), and one due to inaccurate TMS targeting due to a technical error. Additionally, three of these six participants did not complete the task-based fMRI.

The study involved two sessions on separate days. In the first, participants underwent MRI scanning that included structural, resting-state, and task-based fMRI. In the second, they performed a reaching task using the KINARM Exoskeleton Robot (Kinarm, London, ON, Canada) while receiving TMS.

### MRI acquisition

MRI data were collected at the University of Pittsburgh’s Magnetic Resonance Research Center (RRID:SCR_025215) using a 3T Siemens Prisma MRI scanner with a 64-channel head coil.

#### Structural MRI acquisition

Structural images included high-resolution T1-weighted scans acquired using an MP-RAGE sequence (repetition time [TR] = 2300 ms, echo time [TE] = 2.9 ms, inversion time [TI] = 900 ms, flip angle = 9°, voxel size = 1 mm isotropic, acquisition matrix = 256 × 256 × 192). T2-weighted images were collected using a SPACE sequence (TR = 3200 ms, TE = 408 ms, voxel size = 1 mm isotropic, acquisition matrix = 256 × 256 × 192).

#### Resting-state fMRI acquisition

Resting-state fMRI data were acquired while participants had their eyes open, fixating on a central cross displayed on the screen. Imaging parameters were as follows: multiband acceleration factor = 8, TR = 800 ms, TE = 37 ms, voxel size = 2 mm isotropic, acquisition matrix = 104 × 104 × 72, 720 volumes. An additional resting-state scan was collected at the end of the session as a backup in case of data quality issues.

#### Task-based fMRI acquisition

Participants performed a joystick-controlled visuomotor task programmed in E-Prime 2.0 (Psychology Software Tools, Pittsburgh, PA, USA) using their dominant hand. Before the MRI session, participants completed a brief practice session consisting of 24 trials to familiarize themselves with the task. The experiment began with an initial 6-second central fixation. Each trial then started with a fixation period of 2–4 seconds, followed by a 3-second directional cue indicating the target location. Participants moved the joystick cursor horizontally toward a circular target that appeared randomly on the left or right side of the screen. The reaching phase lasted 3 seconds and consisted of approximately 1 second to reach, 1 second to hold, and 1 second to return. A total of 60 trials were presented in randomized order (Fig. 1a). Task-based fMRI data were collected using the same imaging parameters as the resting-state fMRI.

**Fig 1.**
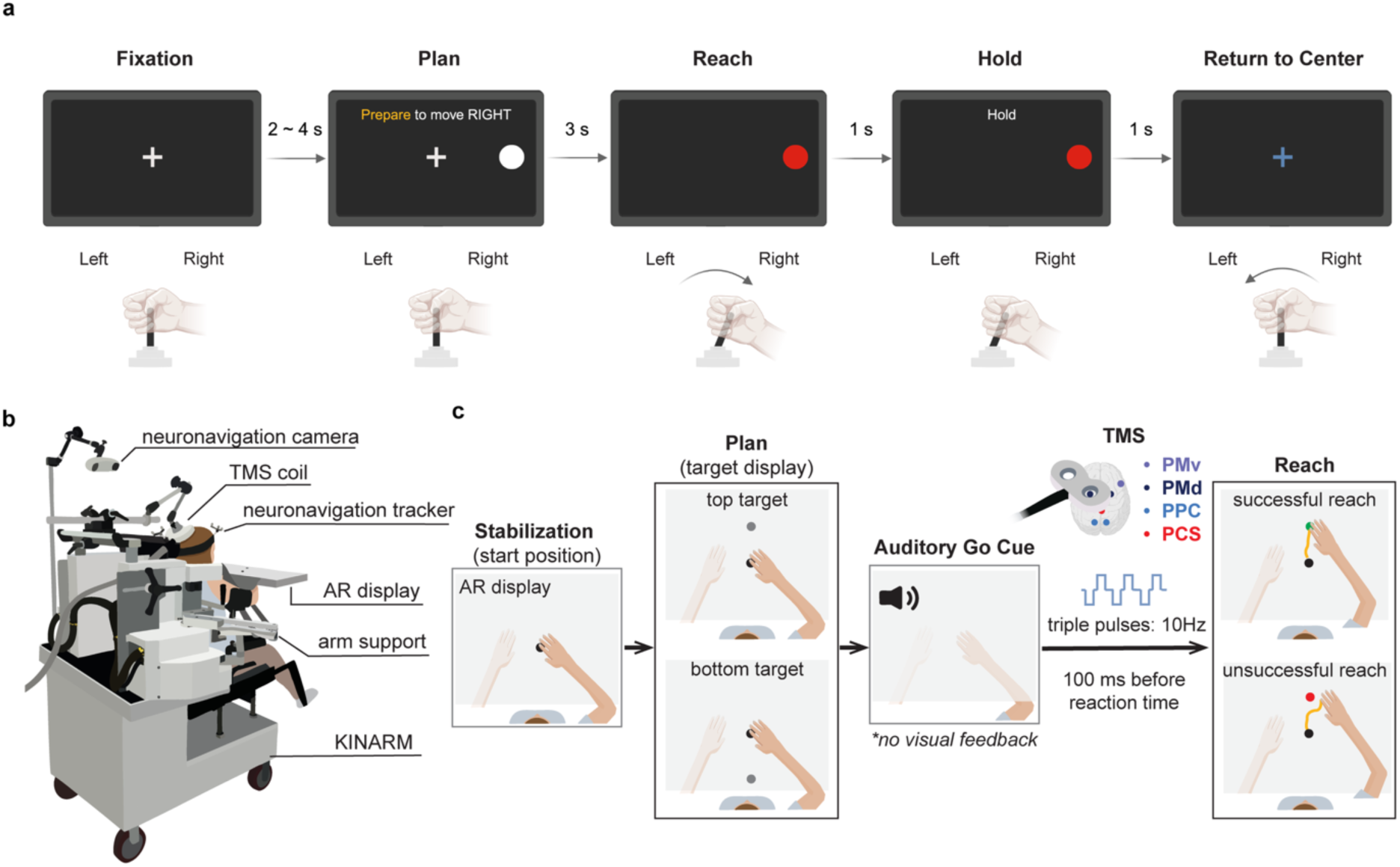
Experimental tasks and setup. **(a)** Schematic of the joystick-controlled visuomotor task performed during fMRI scanning. Each trial included an initial fixation (2–4 s), a planning cue (3 s), and a reaching phase (3 s total: 1 s reach, 1 s hold, 1 s return) toward a left or right target in the horizontal plane. **(b)** Experimental setup integrating the KINARM robotic system with neuronavigated TMS. **(c)** Each trial involved a visual target cue followed by an auditory signal prompting a planar reach. TMS was delivered 100 ms before each participant’s calculated reaction time to bilateral PMd, PMv, or PPC, and PCS was used as the control region. Abbreviations: PMd = dorsal premotor cortex; PMv = ventral premotor cortex; PPC = posterior parietal cortex; PCS = postcentral sulcus.

### Individualized TMS protocol

TMS targets were defined based on cortical areas implicated in reaching control, including bilateral PMd, PMv, PPC, and the postcentral sulcus (PCS) as the control target. Initial coordinates were defined in Montreal Neurological Institute (MNI152; 1-mm isotropic resolution) space using meta-analysis data from NeuroSynth (Yarkoni et al. 2011). The PMv location was based on anatomical landmarks described in previous work (Picard and Strick 2001). All stimulation coordinates are provided in Table 1.

**Table 1.** Coordinates for TMS targets. All coordinates reflect surface-level stimulation targets defined in MNI152 (1-mm) space. Abbreviations: PMd = dorsal premotor cortex; PMv = ventral premotor cortex; PPC = posterior parietal cortex; PCS = postcentral sulcus.

| ROI | Hemisphere | MNI coordinates |  |  |
| --- | --- | --- | --- | --- |
|  |  | x | y | z |
| PMd | Right | 30 | -17 | 71 |
|  | Left | -30 | -17 | 71 |
| PMv | Right | 58 | 6 | 28 |
|  | Left | -58 | 6 | 28 |
| PPC | Right | 21 | -68 | 61 |
|  | Left | -18 | -68 | 61 |
| PCS | Midline | 0 | -60 | 75 |

These coordinates were individualized for each participant by registering the MNI atlas to their T1-weighted MRI using FMRIB’s Linear Image Registration Tool (FLIRT); (Jenkinson and Smith 2001; Jenkinson et al. 2002) from the FMRIB Software Library (FSL; Oxford, UK), coordinates were then transformed into subject space and imported into BrainSight software (Rogue Research, Montréal, QC, Canada) for real-time neuronavigation and accurate coil positioning during the TMS session. Additionally, a 3×3 stimulation grid (10-mm spacing), centered on the anatomical hand knob region of the M1 contralateral to the dominant hand, was defined in BrainSight for motor hotspot mapping and resting motor threshold (RMT) determination. Final stimulation targets were manually adjusted based on individual anatomical variability.

Surface electromyography (EMG) electrodes with integrated amplifiers (B&L Engineering, Santa Ana, CA, USA) were placed over the extensor digitorum communis (EDC) of the dominant arm for motor hotspot identification and RMT determination. Motor hotspot mapping was performed using single-pulse TMS across each of the nine grid locations. The hotspot was identified as the location yielding the largest mean motor-evoked potential (MEP) amplitude and the most consistent MEP responses (≥ 50 µV peak-to-peak amplitude). RMT was then measured at the identified hotspot using the Motor Threshold Assessment Tool (MTAT; Awiszus and Borckardt 2011).

During the reaching task, we applied TMS at three intensities: no-stimulation (0%), subthreshold (80% RMT), and suprathreshold (120% RMT). This range was selected to vary perturbation strength while preserving focality. The subthreshold setting was designed to modulate local networks with limited spread, whereas the suprathreshold setting provided a stronger perturbation to enhance the likelihood of detecting behavioral effects in non-M1 areas, where excitability thresholds cannot be directly measured (Rice et al. 2006). RMT was used to individualize these settings for each participant. For individuals with high RMT values (>83% of maximum stimulator output), suprathreshold stimulation was limited to 100% of maximum output, with subthreshold remaining at 80% RMT.

### KINARM reaching task protocol

Participants were asked to sit upright with their heads comfortably stabilized on a headrest attached to the task screen frame. A PowerMAG PMD70-pCool double coil (Jali Medical, Framingham, MA, USA) was secured on a flexible coil holder attached to the back of the robot, enabling accurate coil positioning and adjustments throughout the experiment (Fig. 1b). Participants performed a planar center-out reaching task using the KINARM Exoskeleton Robot with their dominant hand. Each trial began with the participants stabilizing their fingertip at a central starting position for 1.5 seconds, followed by a visual target presented either forward or backward relative to the center. Following a randomized delay (0.5–1.5 s), an auditory Go cue prompted participants to reach the target as quickly and accurately as possible. Visual feedback of the fingertip (displayed as a white circular cursor) and the target were provided until the Go cue and then removed immediately after (Fig. 1c).

Each participant completed at least 20 practice trials, which were used to calculate reaction time, defined as the latency from the auditory cue to the point where fingertip velocity exceeded 1 cm/s. Kinematic data were recorded at a sampling rate of 1 kHz. The experiment consisted of seven stimulation blocks, one for each of the defined TMS targets. Each block included 36 trials (12 trials per stimulation intensity). Stimulation intensities (no-stimulation, subthreshold, suprathreshold) were randomized within each block, and stimulation targets were randomized using a Latin square design. TMS was delivered as a 10 Hz triple-pulse train initiated 100 ms prior to each participant’s calculated reaction time, specifically targeting the transition from movement planning to execution.

### Data analysis

#### fMRI data preprocessing

To ensure consistent hemispheric alignment across participants, all functional and anatomical images for the two left-handed participants were flipped across the mid-sagittal plane so that the right hemisphere corresponds to the hemisphere ipsilateral to the dominant arm. Resting-state fMRI data preprocessing was performed using the CONN toolbox (Whitfield-Gabrieli and Nieto-Castanon 2012). The pipeline included realignment and unwarping for head motion and distortion correction, segmentation into gray matter, white matter, and cerebrospinal fluid (CSF), spatial normalization to MNI space, and smoothing with a 5-mm Gaussian kernel full width at half maximum (FWHM). Images were resampled to 2-mm isotropic voxels and denoised by regressing out five CompCor components from white matter and CSF (Behzadi et al. 2007), 12 motion parameters with derivatives, and outlier scans (Power et al. 2014), followed by bandpass filtering (0.008–0.09 Hz) to reduce low-frequency drift and high-frequency physiological noise.

Task-based fMRI data were preprocessed using the Statistical Parametric Mapping (SPM12) toolbox (Penny et al. 2011). Preprocessing included spatial realignment to correct for head motion, smoothing with a 4-mm Gaussian kernel (FWHM), and high-pass filtering (0.01 Hz). Anatomical T1-weighted images were automatically segmented using FreeSurfer (Fischl 2012) to improve anatomical localization and enhance the accuracy of functional-to-structural co-registration with each participant’s T1-weighted and T2-weighted images.

#### Resting-State fMRI connectivity analysis

Functional connectivity analysis was based on eight regions of interest (ROIs): bilateral PMd, PMv, PPC, and M1. ROIs were centered on the same coordinates used for stimulation. Each ROI centroid was placed slightly beneath the cortical surface to ensure full gray matter coverage without extending beyond the brain. ROIs were defined as 6.5 mm radius spheres (∼13 voxels), optimized to provide adequate regional coverage while minimizing overlap between adjacent ROIs. Spherical ROIs were created using FSL, merged into a custom atlas, and imported into the CONN toolbox.

ROI-to-ROI functional connectivity was computed in CONN by extracting Blood Oxygen Level Dependent (BOLD) time series as the mean across all voxels within each ROI, followed by calculation of Fisher z-transformed bivariate Pearson correlation coefficients between each ROI pair. At the group level, a general linear model (GLM) was used to test whether each connection was significantly different from zero across participants (one-sample *t*-tests). Connection-level significance was set at *p* < 0.05 and cluster-level inference was based on complete-linkage hierarchical clustering across connections with a false discovery rate (FDR) threshold of *p_FDR_* < 0.05 (Benjamini and Hochberg 1995).

#### Task-based fMRI analysis

Activation maps for the planning and reaching phases were modeled at the single-subject level using a GLM in SPM12. Activity maps were co-registered to each participant’s anatomical scan and normalized to MNI space. For each participant and phase, contrast maps were thresholded using family-wise error (FWE) correction (*p_FWE_* < 0.05, *t* > 5.01). The resulting thresholded maps were then summed voxel-wise across participants to generate group-level overlap maps in MNI space, which were projected onto cortical surface models using AFNI (Analysis of Functional NeuroImages; Cox 1996) and SUMA (Surface Mapping with AFNI; Saad and Reynolds 2012).

#### Kinematic assessment of TMS effects

We analyzed arm kinematics of the planar reaching task to evaluate the effects of different TMS conditions on motor performance, thereby probing the causal relationship between non-primary motor cortical activity and reaching behavior. Hand path trajectories, defined from 1 cm after the start point to 1 cm before the target, were low-pass filtered with a zero-lag, 6^th^-order Butterworth filter at 10 Hz. The resulting trajectories were then quantified using two measures of straightness, path-to-length ratio (PLR) and trajectory deviation error (TDE), and two measures of smoothness, log-dimensionless jerk (LDJ) and spectral arc-length (SAL) (Balasubramanian et al. 2011; Powell et al. 2023). PLR was defined as:

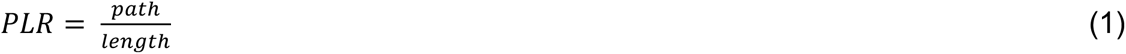

where *path* is the trajectory length, and *length* is the minimum distance between the start position and the end target. TDE was computed as:

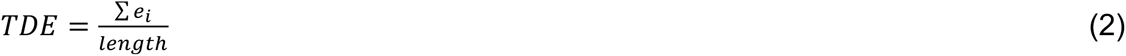

Where *e_i_* is the minimum distance between the *i^th^* sample of the trajectory and the straight line connecting the start point to the end target and *length* is as previously defined. LDJ was used to quantify changes in acceleration:

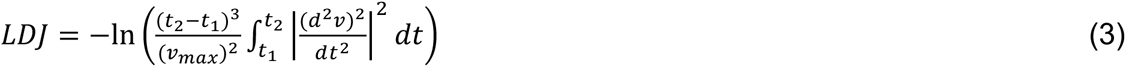

Where *t*_1_ and *t*_2_ represent the start and end times of the trajectory, *v*(*t*) is the hand velocity as a function of time, and *v_max_* is the maximum hand velocity. SAL measured smoothness in the frequency domain:

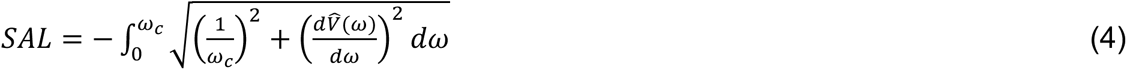

Where *V̂*(*ω*) is the magnitude of the Fourier spectrum of *v*(*t*), normalized by the DC component *V̂*(0), with *ω* as the angular frequency in the range [0, *ω_c_*]. Consistent with previous studies, we adopted *ω_c_* = 40π *rad*/*s* to capture the frequency range characteristic of human movement (McAuley et al. 1997; Balasubramanian et al. 2011). Overall, the straightness measures the geometric deviation of the trajectories from a straight line, whereas the smoothness metrics capture changes in hand acceleration and deceleration during reaching, independent of trajectory geometry.

Trajectory straightness and smoothness metrics were compared independently across the three stimulation conditions (no-stimulation, subthreshold, suprathreshold). To account for inter-individual variability in baseline performance, all metrics were normalized to the values obtained from PCS stimulation. Only trials in which the end target was successfully reached were included, and within-subject condition pairs with fewer than five trials were excluded. We then performed a bootstrap analysis paired within each participant and the reaching target (upper and lower). For each metric, we resampled the paired differences between stimulation target and PCS 10,000 times to generate confidence intervals (CIs). The null hypothesis of no difference in the mean was rejected if zero was not contained within the 95% CI.

#### Functional connectivity patterns associated with reaching performance

To understand how resting-state functional connectivity among cortical areas involved in reaching relates to motor behavior, we assessed associations between connectivity patterns and performance during the reaching task. First, we aimed to determine whether patterns of resting-state functional connectivity, measured by correlations between specific brain regions, were associated with individual differences in baseline reaching performance. Second, we tested whether connectivity patterns could predict changes in motor behavior following TMS applied to different cortical targets and intensities. To address these aims, we used Partial Least Squares Regression (PLSR), a multivariate statistical technique that identifies underlying relationships between functional connectivity measures and behavioral outcomes (Krishnan et al. 2011).

PLSR identifies relationships between two datasets by extracting latent variables from each dataset to maximize covariance between predictors (*X*, connectivity data) and responses (*Y*, behavioral measures). Unlike traditional regression, this method handles multicollinearity and is particularly suitable for high-dimensional datasets common in neuroimaging. PLSR decomposes the predictor and response datasets as:

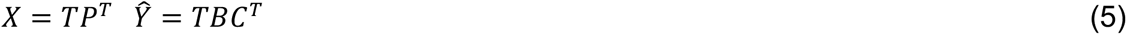

where *T* contains latent variables extracted from *X*, *P* and *C* are loading matrices that capture the contribution of the original variables to the latent variables, and *B* is a diagonal matrix containing regression weights that predict responses from latent variables of *X*.

Here, the predictor variables (*X*) were the average functional connectivity values, z-scored across participants, within each of the nine significant clusters identified from the resting state analysis. The response variables (*Y*) consisted of five behavioral measures from the KINARM reaching task. Three metrics used in the primary KINARM analysis— TDE, PLR, and SAL—represented accuracy, efficiency (trajectory straightness), and smoothness, respectively. We additionally included maximum hand velocity (Max_H_), which reflects peak movement speed, and success rate, defined as the percentage of trials in which the target was reached. Prior to the analysis, we computed pairwise correlations among the behavioral metrics and confirmed that correlations were relatively low (*r* < 0.4), indicating minimal redundancy and suggesting that the metrics measure distinct aspects of performance.

We performed three separate PLSR analyses for each condition. First, we assessed baseline associations between resting-state connectivity (*X*) and behavioral performance (*Y*) by averaging no-stimulation trials across all targets and participants. Next, to evaluate the effects of stimulation, we calculated behavioral change scores for each participant by subtracting their average no-stimulation performance from performance during subthreshold and suprathreshold stimulation conditions, separately for each of the six stimulation targets. These difference scores were then used as response variables in subsequent PLSR analyses, performed separately for each target, with the same connectivity predictors. All behavioral metrics were z-scored prior to analysis. In each analysis, we focused exclusively on the first latent component, which explains the largest shared variance between connectivity and behavior, to simplify interpretation. Statistical significance of this component was assessed using permutation testing with 1000 iterations, and *p*-values were computed as the proportion of permutations showing explained variance equal to or greater than the observed value, with significance set at *p* < 0.05.

## Results

### Cortical activation during planning and reaching

To provide a baseline map of cortical activation during a reaching task in neurologically intact participants for future comparisons with stroke populations, we mapped activation during the planning and reaching phases of the task performed during the fMRI scan (individual maps thresholded at *t* > 5.01, *p_FWE_* < 0.05). All reach directions were pooled across trials. During the planning phase (Fig. 2a), activation was observed in contralateral M1, PMd, and SMA, as well as bilateral PMv, PPC, cerebellum, and primary visual cortex. In the reaching phase (Fig. 2b), activation in PMd, PMv, and PPC remained bilateral and appeared more extensive, with additional engagement of the contralateral sensorimotor cortex and SMA. The cerebellum and visual cortex were consistently active across both phases. Compared to planning, the reaching phase showed broader activation of sensorimotor areas and stronger engagement of bilateral premotor and parietal cortices.

**Fig 2.**
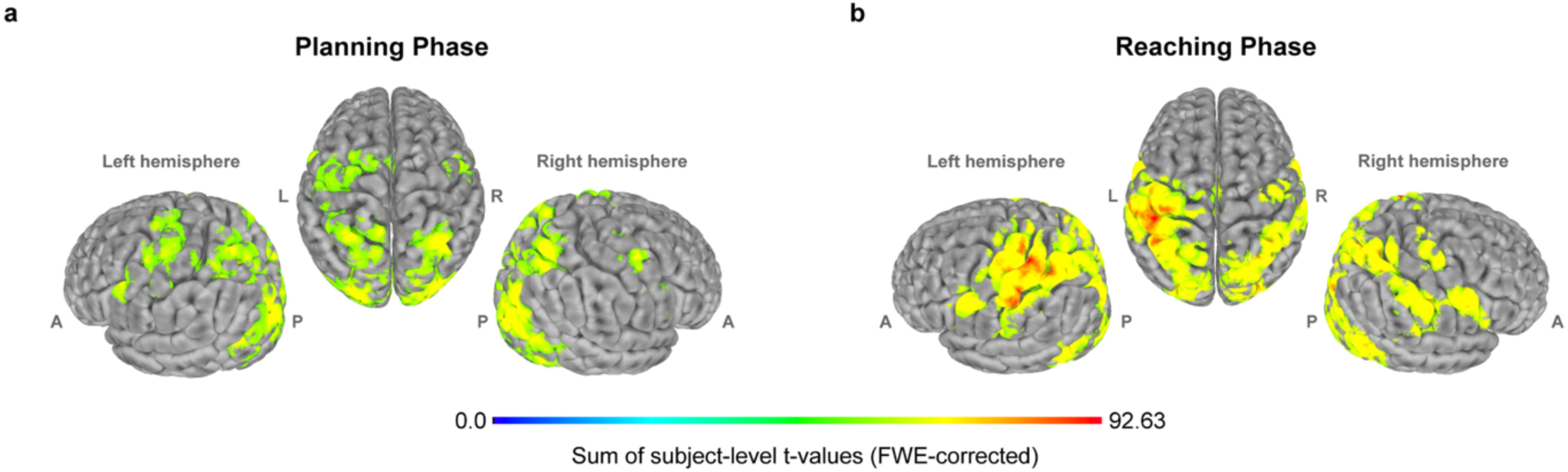
Cortical activation during the reaching task. **(a)** Planning phase. **(b)** Reaching phase. Maps show the voxel-wise sum of subject-level *t*-values from first-level analyses (*p_FWE_* < 0.05, *t* > 5.01) projected onto cortical surfaces. Higher values indicate stronger and more consistent activation across participants. Anatomical directions: anterior (A), posterior (P), left (L), right (R).

### Resting-State connectivity patterns

Group level resting-state connectivity analysis showed significant functional connectivity among our predefined ROIs (bilateral PMd, PMv, PPC, and M1) indicating consistent couplings across participants. Significant ROI-to-ROI connections are illustrated in Fig. 3a. Among the strongest were interhemispheric connections between bilateral PMd (*t* = 16.63), PMv (*t* = 9.96), and M1 (*t* = 9.87). Robust intra- and interhemispheric interactions were also identified between premotor areas (PMd and PMv, *t* = 4.51 to 10.04) and between premotor and primary motor areas (PMd and M1, *t* = 4.68 to 6.18). Additionally, significant interhemispheric connectivity was observed between bilateral PPC (*t* = 5.49). PPC also showed significant connections with premotor areas (*t* = 2.20 to 4.38), along with anticorrelations with M1 (*t* = -2.94 to -4.07). These significant connections were grouped into nine distinct clusters (Table 2), identified through hierarchical clustering based on anatomical proximity and similarity in functional connectivity patterns across participants. Clusters were identified using a connection-level threshold of *p* < 0.05 and cluster-level threshold of *p_FDR_*< 0.05.

**Fig 3.**
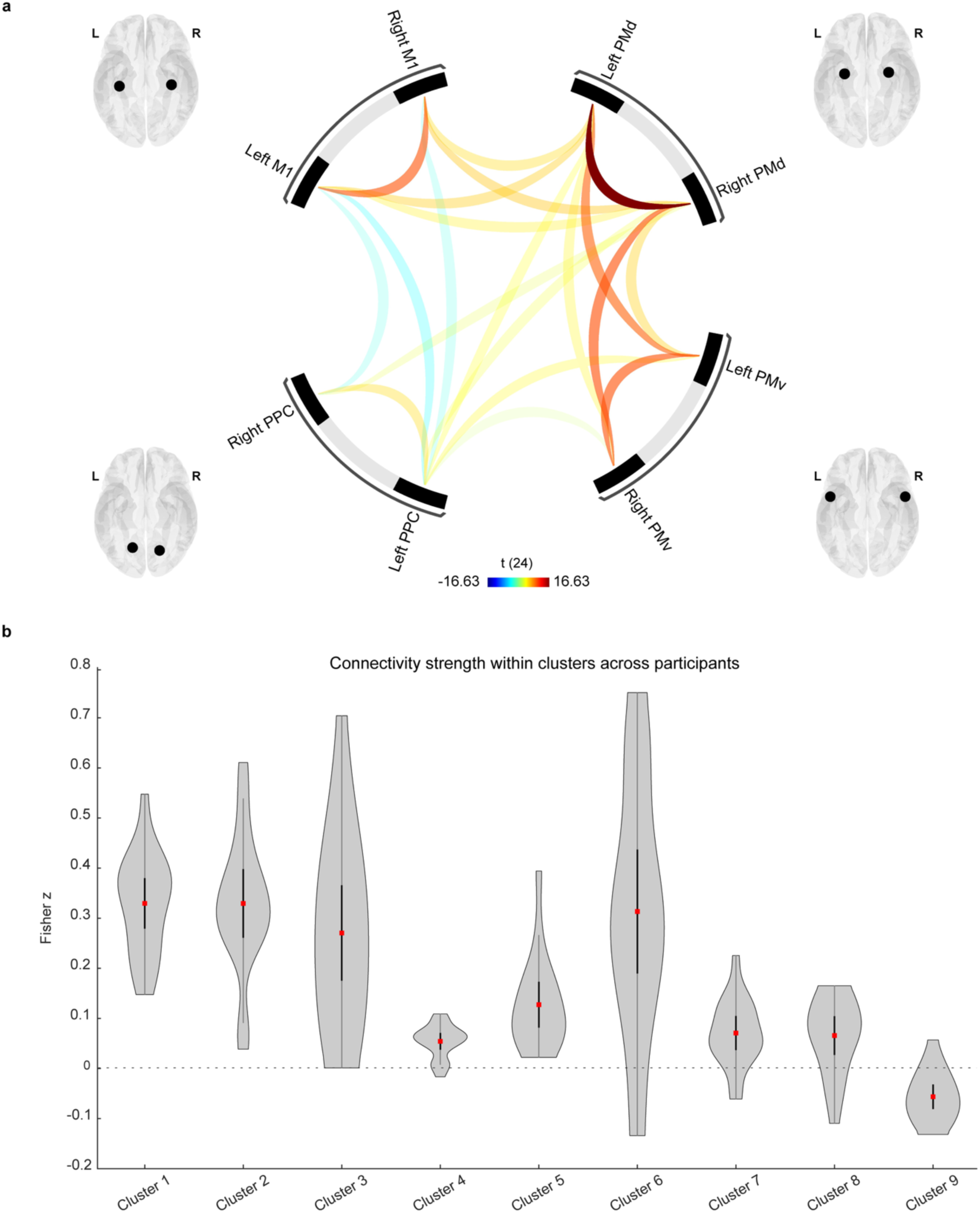
Resting-state functional connectivity among ROIs. **(a)** Significant ROI-to-ROI connections identified at the group level (*p_FDR_* < 0.05). Warm colors indicate positive connectivity (*t* > 0), and cool colors indicate anticorrelations (*t* < 0). Line thickness reflects *t*-value magnitude. **(b)** Violin plots display the distribution of Fisher z-transformed connectivity values for each of the nine clusters across participants. Red squares indicate the mean, and error bars represent the 95% CI. Abbreviations: PMd = dorsal premotor cortex; PMv = ventral premotor cortex; PPC = posterior parietal cortex; M1 = primary motor cortex.

**Table 2.**
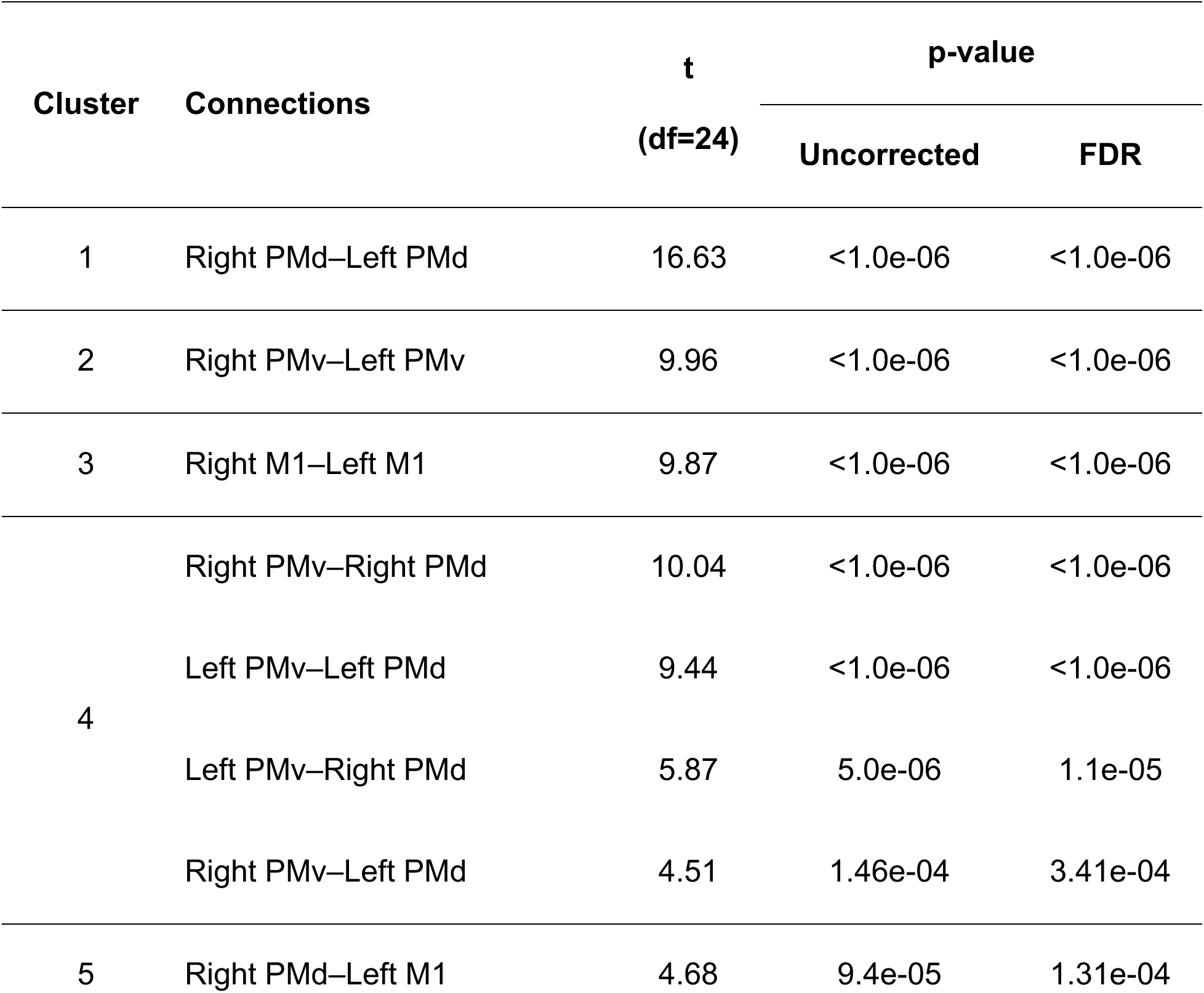
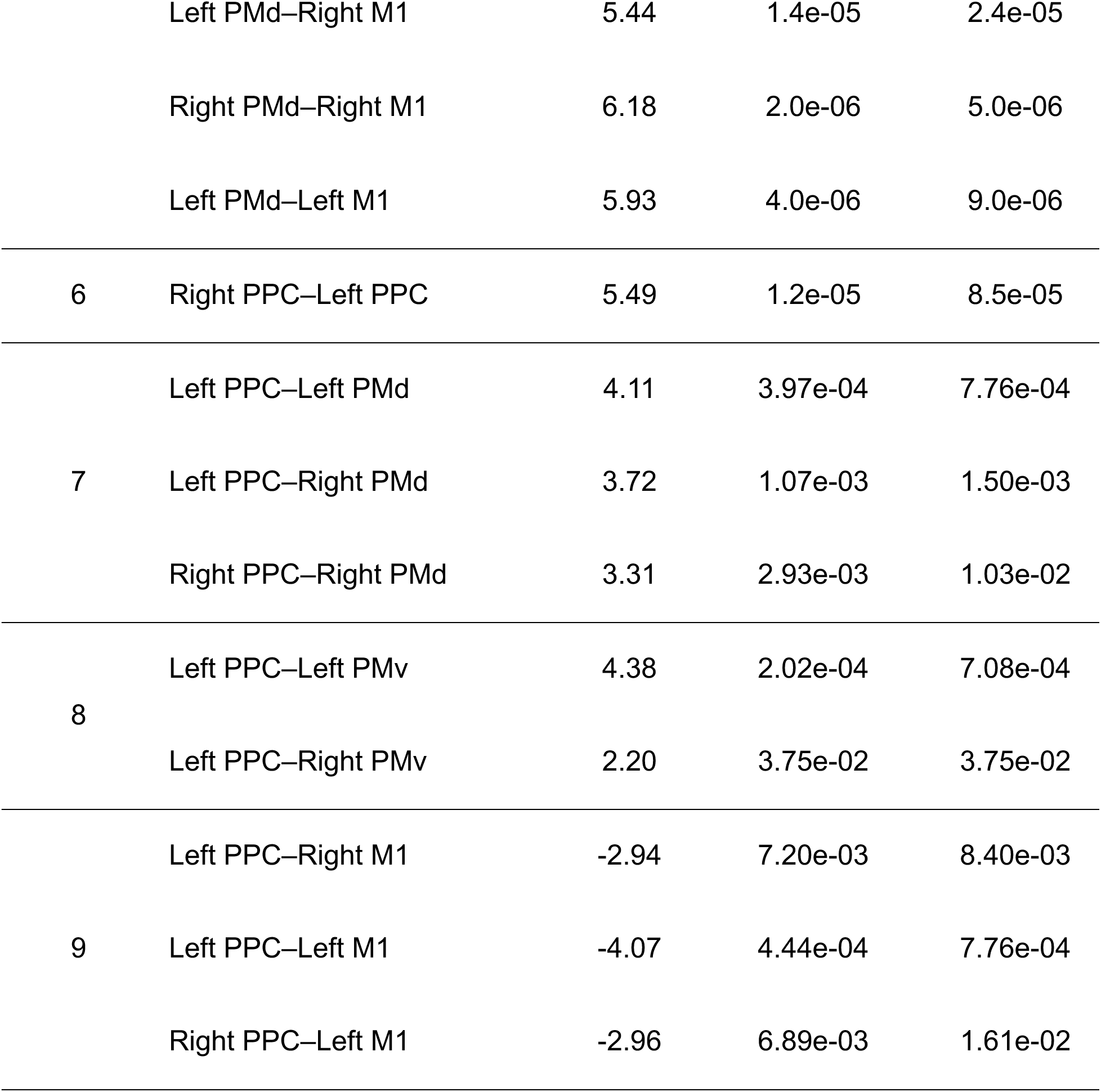
Significant functional connectivity clusters. Each cluster shows the connections between ROIs, corresponding *t*-values (df = 24), and associated *p*-values (uncorrected and FDR-corrected). Abbreviations: PMd = dorsal premotor cortex; PMv = ventral premotor cortex; PPC = posterior parietal cortex; M1 = primary motor cortex.

For subsequent analyses, Fisher z-transformed connectivity values were averaged for each participant across the ROI-to-ROI connections that defined each cluster. The distributions of these cluster-wise averages are shown in Fig. 3b. Across participants, interhemispheric premotor clusters (Clusters 1 and 2) showed consistently positive connectivity (Cluster 1: mean = 0.33 ± 0.10; Cluster 2: mean = 0.33 ± 0.14). Among clusters, Interhemispheric M1 (Cluster 3: mean = 0.27 ± 0.20) and bilateral PPC (Cluster 6: mean = 0.31 ± 0.25) displayed the highest interindividual variability. Smaller clusters, including intra- and interhemispheric premotor couplings (Cluster 4: mean = 0.05 ± 0.03) and PPC–premotor clusters (Clusters 7 and 8: means ≈ 0.07 ± 0.07), showed weaker but reliable positive connectivity across subjects. Finally, PPC– M1 connections (Cluster 9) were anticorrelated (mean = -0.06 ± 0.05) across participants. The clusters were used to group anatomically and functionally related connections into compact, interpretable modules. This reduced dimensionality while preserving the overall network structure, facilitated the assessment of between-subject variability, and provided a more interpretable structure for analyzing connectivity– behavior relationships.

### Effects of TMS on reach trajectories

We assessed how TMS of non-primary motor areas during movement planning, delivered 100 ms before each participant’s reaction time (mean = 339.71 ± 81.35 ms, range = 233–515 ms), affected reach trajectories by comparing trajectory straightness and smoothness metrics. In each trial, participants stabilized their hand at the start position before reaching an upper or lower target on the AR display (Fig. 4a). We compared the effects of TMS across three stimulation conditions defined relative to each participant’s RMT (mean = 74.29% of maximum stimulator output ± 13.18, range = 50–100% of maximum stimulator output). These included no-stimulation, subthreshold (80% RMT), and suprathreshold (120% RMT).

**Fig 4.**
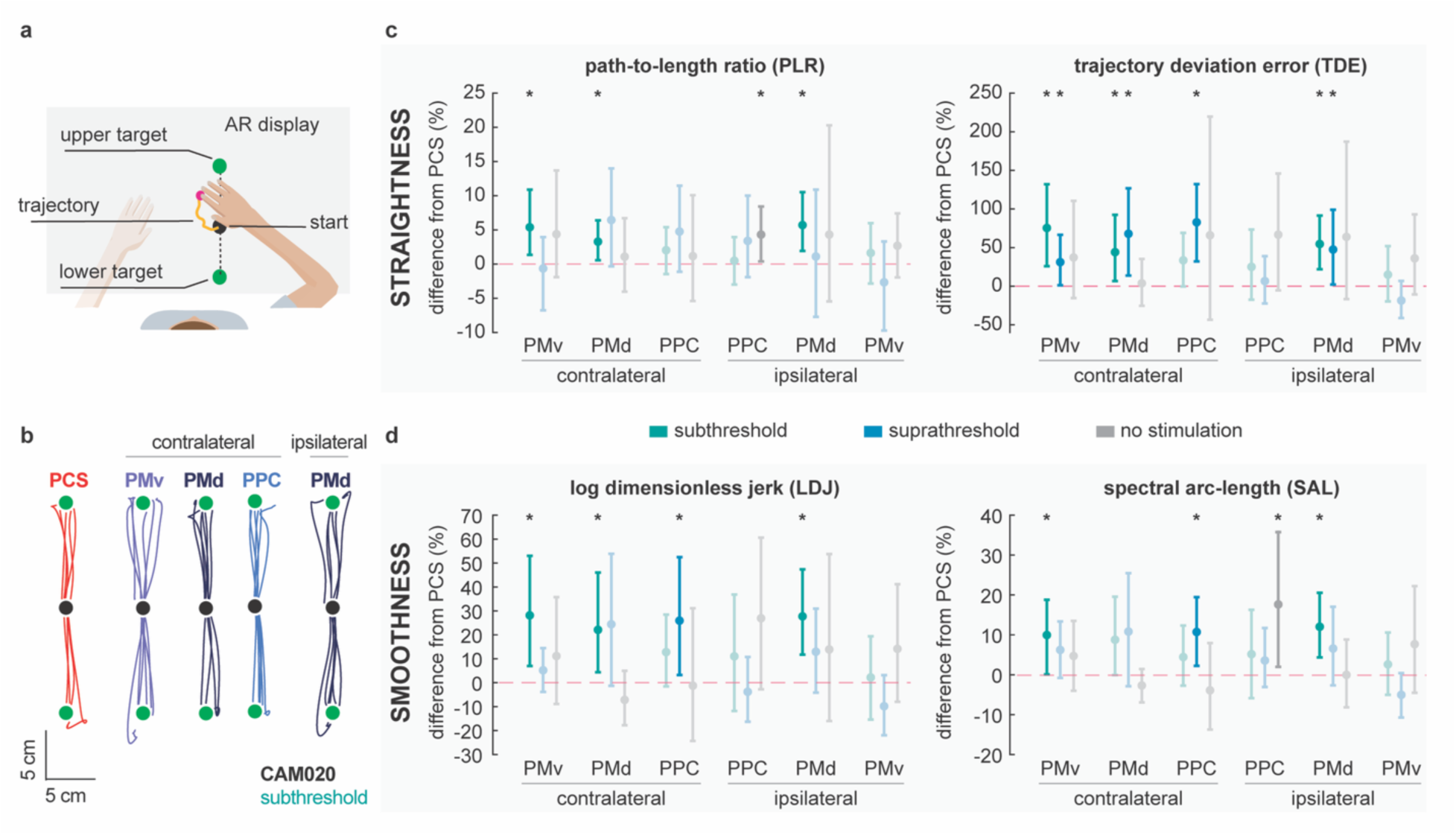
TMS-induced changes in reach trajectories during the KINARM task. **(a)** Layout of the start position and target locations (upper and lower) displayed during the reaching task. **(b)** Example trajectories from a participant following stimulation of contralateral PMv, PMd, and PPC and ipsilateral PMd, relative to PCS (control target). **(c)** Percent change in trajectory straightness (PLR and TDE) relative to PCS across stimulation targets. **(d)** Percent change in trajectory smoothness (LDJ and SAL) relative to PCS. Subthreshold, suprathreshold, and no-stimulation conditions are shown in green, blue, and gray, respectively. Asterisks indicate significant differences relative to PCS (bootstrap, 95% CI). Abbreviations: PMd = dorsal premotor cortex; PMv = ventral premotor cortex; PPC = posterior parietal cortex; PCS = postcentral sulcus.

In Fig. 4b, example trajectories from one participant illustrate that subthreshold stimulation of contralateral PMv, PMd, and PPC, as well as ipsilateral PMd, resulted in more curved and variable reach paths compared to the PCS control target. These disruptions were consistent across participants and confirmed by quantitative analyses.

We found that reach trajectories were less straight and smooth during subthreshold TMS of contralateral PMv and bilateral PMd compared to PCS (Fig. 4c, d). These effects were reflected in higher TDE/PLR and increased LDJ and SAL. Although subthreshold TMS of contralateral PPC did not seem to affect reaching movements, suprathreshold stimulation of contralateral PPC significantly reduced both straightness and smoothness measures. For all these stimulation sites, trials without any stimulation were not significantly different from the control target, confirming that the observed disruptions were influenced by TMS during movement planning. Interestingly, reach trajectories were affected in the no-stimulation condition when the coil was positioned over ipsilateral PPC, possibly reflecting a side effect of coil placement relative to the control target (PCS), as the coil remained on the scalp at 0% output during all no-stimulation trials. Overall, results show that stimulation of contralateral PMv, PMd, PPC, and ipsilateral PMd influences the straightness and smoothness of reach trajectories, and among stimulation intensities, subthreshold stimulation led to the most consistent effects, likely due to its more local influence without engaging broader regions.

### Resting-State connectivity predictors of reaching performance

We used PLSR to assess whether resting-state functional connectivity among motor-related brain areas could predict individual differences in reaching behavior, both in the absence of stimulation and following TMS delivered to different cortical targets and at different intensities. The predictors (*X*) consisted of Fisher z-transformed connectivity values from nine ROI-to-ROI clusters, and the response variables (*Y*) included five behavioral metrics derived from the KINARM task.

In the no-stimulation condition, the first PLS component explained 11.7% of the variance in behavior and 39.9% of the variance in connectivity, but the model was not statistically significant (*p* = 0.661). Among stimulation conditions, a significant effect on reaching behavior was observed only with subthreshold TMS of the right PMd (ipsilateral to the dominant arm), which explained 27.6% of the variance in behavior and 35% of the variance in connectivity (*p* = 0.034; permutation test in Fig. 5a, d). Suprathreshold stimulation of right PMd also showed a trend (*p* = 0.066). All other stimulation targets, including those in the left hemisphere, showed no significant associations with resting-state connectivity values (*p* > 0.35). The connectivity loadings (Fig. 5b) identified cluster 3 (bilateral M1) as the strongest positive contributor and cluster 6 (bilateral PPC) as the strongest negative. The behavioral loadings (Fig. 5c) showed that TDE had the highest positive weight, with smaller negative contributions from SAL, Max_H_, and success rate. This pattern indicates that participants with stronger bilateral M1 connectivity and weaker PPC connectivity reflected stronger changes and disruptions in spatial accuracy following right PMd stimulation.

**Fig 5.**
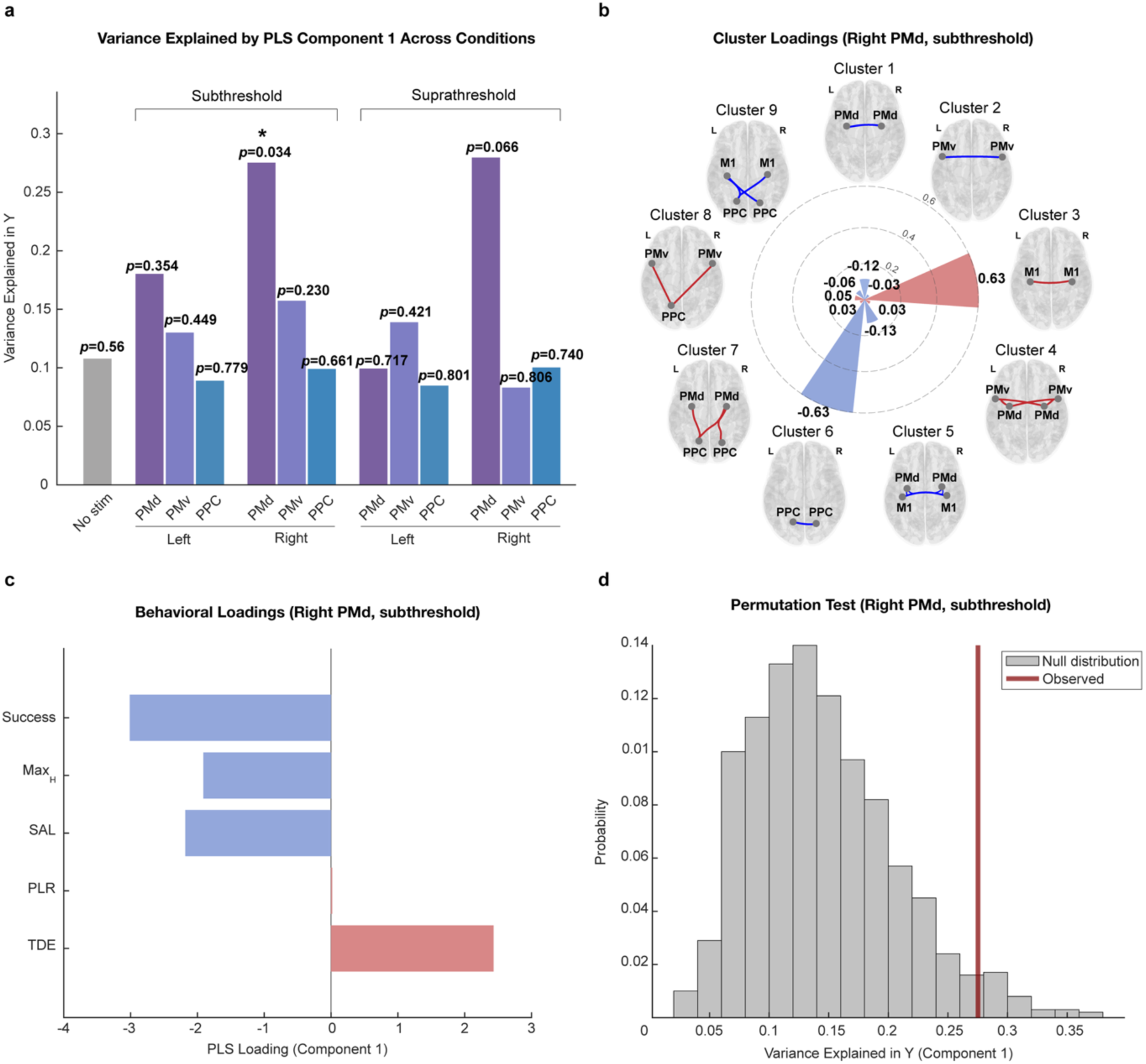
Partial Least Squares (PLS) analysis of connectivity-behavior relationships across all stimulation conditions. **(a)** Explained variance in behavioral data (Y) for each stimulation target and intensity. **(b)** Connectivity cluster loadings for the significant right PMd (ipsilateral to the dominant arm) subthreshold condition. Arc length indicates loading magnitude; red arcs are positive, and blue arcs are negative. Corresponding brain areas for each cluster are shown (cluster colors illustrate the loading direction). **(c)** Behavioral loadings for the right PMd subthreshold condition. **(d)** Permutation test comparing observed explained variance (red line) with the null distribution (gray bars) for the significant condition. Abbreviations: PMd = dorsal premotor cortex; PMv = ventral premotor cortex; PPC = posterior parietal cortex; Max_H_ = maximum hand velocity; SAL = spectral arc-length; PLR = path-to-length ration; TDE = trajectory deviation error.

## Discussion

In this study, we used an approach that involved both measuring and perturbing brain activity, as well as behavior, to investigate how non-primary motor areas shape movement planning and execution in reaching performance. While previous studies have applied TMS to study the cortical contributions to movement planning, our integration of a robotic environment enabled precise quantification of movement features such as smoothness, accuracy, and straightness, which cannot be captured by conventional behavioral or kinematic assessments. Using this approach, we observed that stimulation of contralateral PMd, PMv, PPC, and ipsilateral PMd during the planning phase led to reductions in smoothness and straightness of reach trajectories.

Additionally, we tested whether individual differences in resting-state connectivity could explain behavioral responses to stimulation across all cortical targets. Among all cortical targets, connectivity predicted behavioral responses only for subthreshold stimulation of right PMd (ipsilateral to the dominant arm), where stronger interhemispheric M1 connectivity and weaker PPC connectivity were associated with greater TMS-induced effects.

To establish a baseline map of cortical areas engaged in movement planning and execution for later comparisons with a stroke cohort, we mapped brain activation during a joystick-controlled reaching task using fMRI. We observed significant activation of PMd, PMv, PPC, SMA, and the cerebellum during both the planning and reaching phases of the task. This aligns with prior work by Bartolo (Bartolo et al. 2014), which showed that deciding whether a target is within reach activates similar frontoparietal and cerebellar areas, even without any actual movement. We then applied TMS to the same accessible cortical areas, bilateral PMd, PMv, and PPC, to causally probe their contributions to specific components of reaching.

Stimulation of contralateral PMd, consistently disrupted movement efficiency, spatial accuracy, and smoothness, particularly at subthreshold intensity. This aligns with an earlier study showing that PMd influences M1 activity during movement preparation (Koch et al. 2006), and that disrupting PMd interferes with trajectory formation and anticipatory motor control (Chouinard and Paus 2006). O’Shea (O’Shea et al. 2007) further demonstrated that PMd–M1 connectivity modulates reaction timing depending on the effector and task demands. Similarly, stimulation of contralateral PMv led to disruptions in accuracy, efficiency and smoothness. While previous work has shown PMv’s role in grasp control, particularly in finger positioning and sequencing during precision tasks (Davare et al. 2006), other studies have pointed to its broader involvement in object-directed action and its integration within parieto-frontal networks (Grol et al. 2007). Gallivan (Gallivan et al. 2011) further showed that PMv encodes hand-specific motor plans even before movement execution. Our results extend these findings by demonstrating that PMv also plays a causal role in guiding reach-related movements.

In addition to the frontal premotor areas, PPC contributed to the initial movement planning. Prior work by Desmurget (Desmurget et al. 1999) indicated that PPC plays a key role in updating ongoing reaching movements, and applying TMS over PPC disrupts error correction during goal-directed actions. Consistent with this, we found that suprathreshold stimulation of contralateral PPC increased deviation error and reduced movement smoothness. This aligns with findings from (Glover et al. 2005), where rTMS over the intraparietal sulcus impaired online adjustments to object size. Additionally, Rushworth (Rushworth et al. 2001) showed that the left PPC helps direct spatial attention during hand movements. This suggests that stimulating PPC may also interfere with the attention needed to guide the hand accurately.

One of the main strengths of this study is the integration of resting-state connectivity with behavioral effects of TMS. Using PLSR, we assessed whether individual differences in connectivity across motor-related cortical areas could explain variation in TMS responses. Among targets ipsilateral to the arm being moved, only PMd showed consistent disruptive effects on both straightness and smoothness metrics. This pattern was further supported by PLSR results, which identified the right PMd as the only stimulation target where resting-state connectivity significantly predicted behavioral changes. Interestingly, the predictive connections involved bilateral M1 and PPC regions, rather than PMd connections. This pattern is consistent with prior work showing that stimulation effects can be predicted by the connectivity of remote regions, even when the stimulated site itself is not involved in the predictive network (Gießing et al. 2020). Participants with stronger bilateral M1 connectivity and weaker bilateral PPC connectivity were more likely to show greater TMS-induced changes in spatial accuracy and path efficiency. These findings suggest that the behavioral impact of stimulating PMd may depend on the broader interhemispheric motor and parietal network organization. Comparing functional connectivity across motor networks in stroke patients to that of individuals with intact networks may help identify compensatory or maladaptive patterns. Future studies could build on this approach by assessing stimulation responsiveness based on baseline connectivity in clinical populations.

Several limitations should be acknowledged in this study. First, the sample size was modest, which may limit statistical power and generalizability. Second, although we targeted accessible cortical areas, deeper or less accessible areas such as SMA or cerebellum were not directly investigated at this stage. Additionally, stimulation timing was based on reaction times measured during initial practice, which may not reflect changes due to fatigue over the course of the experiment. Another limitation concerns the task design. In the fMRI component, the joystick task was mainly performed by wrist movements, yet it still required goal-directed actions along defined trajectories that capture key aspects of reaching. The KINARM task was also simplified to planar movements, which may limit how well our findings generalize to more natural, unconstrained reaching. Still, Urbin (Urbin et al. 2021) reported that perturbing premotor areas also altered trajectory curvature and variability in unconstrained reaches, suggesting our results are likely relevant beyond the simplified task. Future studies should explore more diverse movement directions, adaptive stimulation timing, and additional cortical targets.

In conclusion, our results show that non-primary motor areas contribute to reach control, and that the behavioral impact of PMd stimulation in neurologically intact individuals is associated with individual differences in the connectivity of broader motor and parietal networks. Extending this approach to stroke populations may help identify predictive connectivity patterns that inform the selection of neuromodulation targets. By comparing disrupted networks in stroke with those of intact individuals, future studies can work toward more precise and personalized intervention strategies.

## Acknowledgments

We would like to thank Tyler Madonna and Tyler Simpson for technical support with the KINARM robot and software. We also thank Debbie Harrington and Sydney Bader for their help with participant recruitment. We are grateful to Dr. Peter Strick for a discussion regarding the targeting of the premotor region—PMv in particular—and to Dr. Mike Urbin for valuable discussions on the study design and interpretation.

Corresponding author: Golnaz Haddadshargh; University of Pittsburgh, Rehab Neural Engineering Labs, 1622 Locust St, 4th Floor, Pittsburgh, PA 15219.

## Funding

This work was supported by the Veterans Health Administration Rehabilitation Research. & Development Program Merit Review Award #1I01RX003511-01A2. The authors declare that there are no conflicts of interest.

## Notes

### Competing Interest Statement

The authors have declared no competing interest.

